# Individualized connectome-targeted transcranial magnetic stimulation for neuropsychiatric sequelae of repetitive traumatic brain injury in a retired NFL player

**DOI:** 10.1101/151696

**Authors:** Shan H. Siddiqi, Nicholas T. Trapp, Pashtun Shahim, Carl D. Hacker, Timothy O. Laumann, Sridhar Kandala, Alexandre R. Carter, David L. Brody

## Abstract

The recent advent of individualized resting-state network mapping (RSNM) has revealed substantial inter-individual variability in anatomical localization of brain networks identified using resting-state functional MRI (rsfMRI). Such variability may be particularly important after repetitive traumatic brain injury (TBI), which is associated with treatment-resistant depression. RSNM enables personalized targeting of repetitive transcranial magnetic stimulation (rTMS), a focal brain stimulation technique that relieves depression when administered over dorsolateral prefrontal cortex.

RSNM was used to identify left/right dorsolateral prefrontal rTMS targets with maximal difference between dorsal attention network and default mode network (DMN) correlations. These targets were spatially distinct from those identified by prior methods. The method was evaluated by administering twenty sessions of left-sided excitatory and right-sided inhibitory rTMS to a retired NFL defensive lineman with progressive treatment-resistant neuropsychiatric disturbances. Treatment led to improvement in Montgomery-Asberg Depression Rating Scale (72%), cognitive testing, and headache scales. In comparison with healthy individuals and subjects with TBI-associated depression, baseline rsfMRI revealed substantially elevated DMN connectivity with medial temporal lobe (MTL). Serial rsfMRI scans showed gradual improvement in MTL-DMN connectivity and stimulation site connectivity with subgenual anterior cingulate cortex. This highlights the possibility of individualized neuromodulation and biomarker-based monitoring for neuropsychiatric sequelae of repetitive TBI.

## Introduction

Repetitive transcranial magnetic stimulation (rTMS) is a neuromodulatory technique with antidepressant^1,2^ and neurorehabilitative effects^3^. rTMS selectively modulates cortical excitability^4^, which is often affected in traumatic brain injury (TBI)^5^. rTMS has thus been proposed as a potential treatment for patients with TBI-associated depression^6^, especially given that these patients may be less responsive to antidepressant pharmacotherapy^7,8^. While concern for rTMS-induced seizure risk often limits its use in TBI, this risk appears to be elevated primarily in penetrating/hemorrhagic injuries rather than diffuse/multifocal axonal injury^9,10^.

rTMS for major depressive disorder is traditionally targeted to scalp regions approximately overlying dorsolateral prefrontal cortex (DLPFC)^11^, which likely modulates activity in the subgenual anterior cingulate cortex (sgACC)^12^. This concept dates back to the successful use of rostral cingulotomy, a neurosurgical procedure which was first used in the 1950s to disrupt connectivity between prefrontal cortex and deep limbic regions^13^. Subsequent neuroimaging studies confirmed that depression is associated with hyperactivity in sgACC and anti-correlated hypoactivity in DLPFC^14^. This sgACC hyperactivity appears to be normalized when antidepressant pharmacotherapy^15^ and electroconvulsive therapy^16^ are effective, but not when these treatments are ineffective. In light of these findings, deep brain stimulation (DBS) has successfully been used to treat depression by directly attenuating sgACC hyperactivity^17^.

Similar principles have also been proposed for identifying rTMS targets. Across a wide variety of neuropsychiatric disorders, functional connectivity (FC) studies show that optimal targets for excitatory rTMS appear to be functionally anti-correlated with optimal DBS targets for the same disorder^2^. In the case of major depression, rTMS responders have stronger DLPFC target anti-correlation with subgenual anterior cingulate cortex (sgACC)^18^ and stronger sgACC correlation with default mode network (DMN) than treatment non-responders^12^.

While FC-based targeting of regions anti-correlated with sgACC has demonstrated promise in large-group studies, this approach has so far been unsuccessful for identifying patient-specific treatment targets^2,18,19^. This may be explained by inter-individual variability in functional localization of resting state networks (RSNs), a series of large-scale brain systems composed of various anatomical regions that coordinate to perform specific functions^20,21^. Such variability in RSN anatomy is not reliably identified by FC with seeds derived from group averages^22^. The recent advent of iterative correlation and classification algorithms, which can reliably partition a human brain into several individualized RSN maps^22-25^, may therefore enable development of patient and disease-specific rTMS targeting.

Because recent cortical topographic maps classify sgACC as part of DMN^26^, mapping this network may serve as a reasonable method for determining individualized sgACC connectivity profiles. DMN is most strongly anti-correlated with dorsal attention network (DAN), which contains dorsolateral prefrontal nodes that show substantial inter-individual topographic variability^21^. In major depression, these dorsolateral prefrontal regions show hypoconnectivity with DAN and hyperconnectivity with DMN^27^. DBS of sgACC has been shown to modulate interactions between attention-switching and self-referential emotional engagement, functions that are mediated by DAN and DMN, respectively^28^. Due to these parallels with the aforementioned DLPFC-sgACC anti-correlation in major depression, mapping DANDMN anti-correlation may serve as a reasonable proxy for identifying individualized rTMS targets that will modulate DLPFC-sgACC interactions.

Individualized functional localization is particularly challenging in traumatic brain injury (TBI), which causes multifocal white matter injury^29^ and less-predictable FC changes^30^. However, there is a need for neuromodulatory treatment approaches given the lack of effective antidepressant treatments for TBI-associated depression^8^. Like major depression, TBI can affect FC in anterior cingulate cortex (ACC), DLPFC, DAN, and DMN^31-35^. Repetitive TBI has additionally been associated with neurodegenerative changes in medial temporal lobe (MTL) on histopathology and positron emission tomography^36,37^. Due to these changes, patients with repetitive head trauma provide a unique opportunity to investigate not only the nature of the circuitry underlying its neuropsychiatric phenotype, but also into the nature of brain network dynamics in response to noninvasive brain stimulation. Thus, we hypothesized that stimulating the DLPFC target with maximal subject-specific DAN-DMN anti-correlation would modulate sgACC and improve mood in repetitive TBI.

## Methods

### Standard protocol approvals, registrations, and participant consents

This study was conducted in accordance with a protocol approved by the Human Research Protection Office at Washington University School of Medicine in St. Louis. All individuals gave informed written consent. The study was registered with ClinicalTrials.gov (NCT02980484). The study was reviewed regularly by the investigators for safety.

### Subjects

#### Patient

We evaluated a man in his fourth decade of life with a history of neuropsychiatric illness associated with repetitive head trauma during a prior career as a defensive lineman in the National Football League (NFL) and at the amateur level. He reported a history of at least 12 prior concussions, including at least two at the amateur level and at least 10 in the NFL. He likely experienced at least 7,000 sub-concussive head impacts prior to his NFL career, as estimated by the Cumulative Head Impact Index^38^, and an unknown but likely comparable number during his NFL career. He described a 2-3 year history of progressively worsening depression, anxiety, impulsivity, anger, and neurocognitive impairment (particularly long-term and short-term memory). He was unable to work and had restricted social function because of cognitive impairments and emotional dysregulation. He had previously demonstrated inadequate response to sertraline, paroxetine, and alprazolam and was not taking any neuropsychiatric medications at the time of the study. His conventional MRI scan was within normal limits (figure S1). He was enrolled from the Washington University TBI clinic as part of a pilot doubleblind randomized-controlled trial of rTMS for depression associated with TBI with a planned sample size of 20.

##### Controls with TBI-associated depression

10 additional subjects with a history of depression and TBI (8 males, ages 19 to 64) received rsfMRI scans as part of the aforementioned randomized-controlled trial with the same imaging protocol as the experimental subject. This included patients with a score of at least 10 on the Montgomery-Asberg Depression Rating Scale (MADRS) and a history of one to two TBIs associated with low risk of seizure disorder, but no history of repetitive concussive or subconcussive events.

##### Healthy Controls

The healthy control group included 12 male volunteers (ages 30 to 36) with no reported TBI history who received rsfMRI scans as part of the Human Connectome Project (HCP)^39^.

### Clinical Assessments

Clinical testing at baseline and after the full course of treatment included depression testing with MADRS; personality testing with the temperament and character inventory (TCI); self-report mood scales in the NIH Toolbox Emotion Battery (EB) and TBI Quality of Life scale (TBI-QoL); cognitive testing with the NIH Toolbox Cognitive Battery (CB); self-report headache Likert scores and six-question Headache Impact Test (HIT-6); and an expert psychiatric evaluation based on DSM-5 diagnostic criteria. Structural and functional MRI scans were performed at baseline and at the end of the treatment course. Unblinded MADRS, TCI, and EB were repeated at a follow-up assessment six weeks after the completion of the treatment course; CB and TBI-QoL were not repeated due to the subject’s preference. MADRS was the primary outcome measure for the double-blind randomized-controlled trial.

### MRI acquisition and analysis

Full methodological details are presented in the supplement. Pre- and post-treatment MRI included one T1-weighted sequence and 16.5 minutes of blood oxygen level dependent (BOLD) fMRI sequences. Data were pre-processed following Power et al., 2014^40^ (figure S2) and individualized RSN estimation was conducted following Hacker et al., 2013 to determine each voxel’s likelihood of membership in each of seven RSNs^22^. The resulting individualized RSN maps were used to identify TMS targets at left- and right-sided voxel clusters in DLPFC with maximal difference between DAN and DMN (figure 1). Alternate targets were determined based on previously-reported methods based on anatomical landmarks, structural MRI, and rsfMRI.

**Figure 1.**
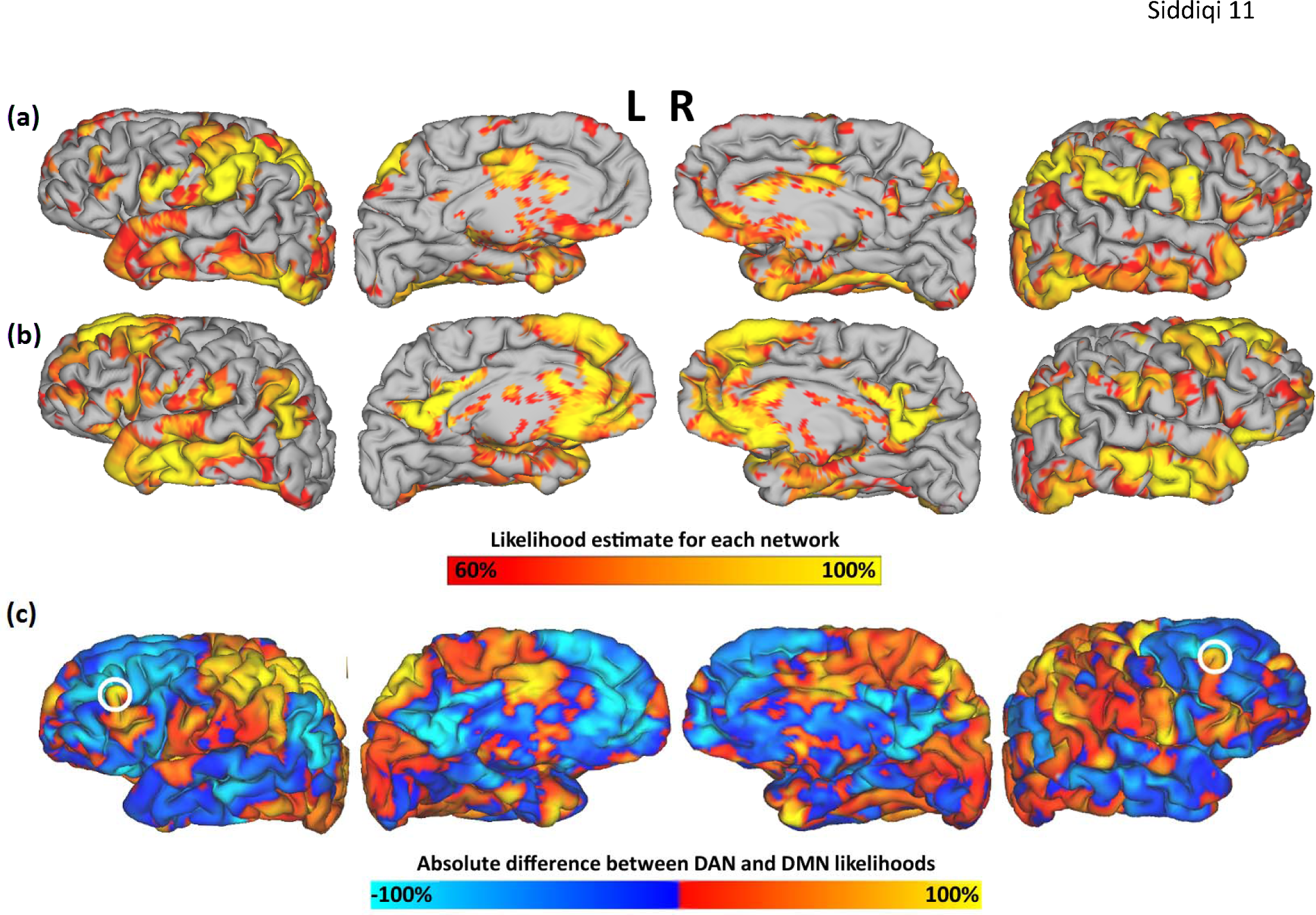
Individualized maps of voxel-wise likelihood estimates for dorsal attention network (a) and default mode network (b) thresholded to display voxels with at least 60% likelihood and visualized as surface projections on a three-dimensional reconstruction of the subject’s anatomical MRI scan. (c) Absolute difference between dorsal attention and default mode network likelihood. Circled regions reflect the TMS stimulation sites, chosen to be the dorsolateral prefrontal regions with maximal difference between DAN and DMN correlations, as determined by subtracting (b) from (a).

A whole-brain “winner take all” parcellated map was generated by assigning each voxel to the network at which it demonstrated the highest likelihood of membership. These maps were used to subjectively compare RSN topography between the experimental subject, the group of healthy controls, and a representative example of a healthy control (figure 2). The seven networks identified by this parcellated map were used as regions of interest (ROIs) for seed-based connectivity analysis.

**Figure 2.**
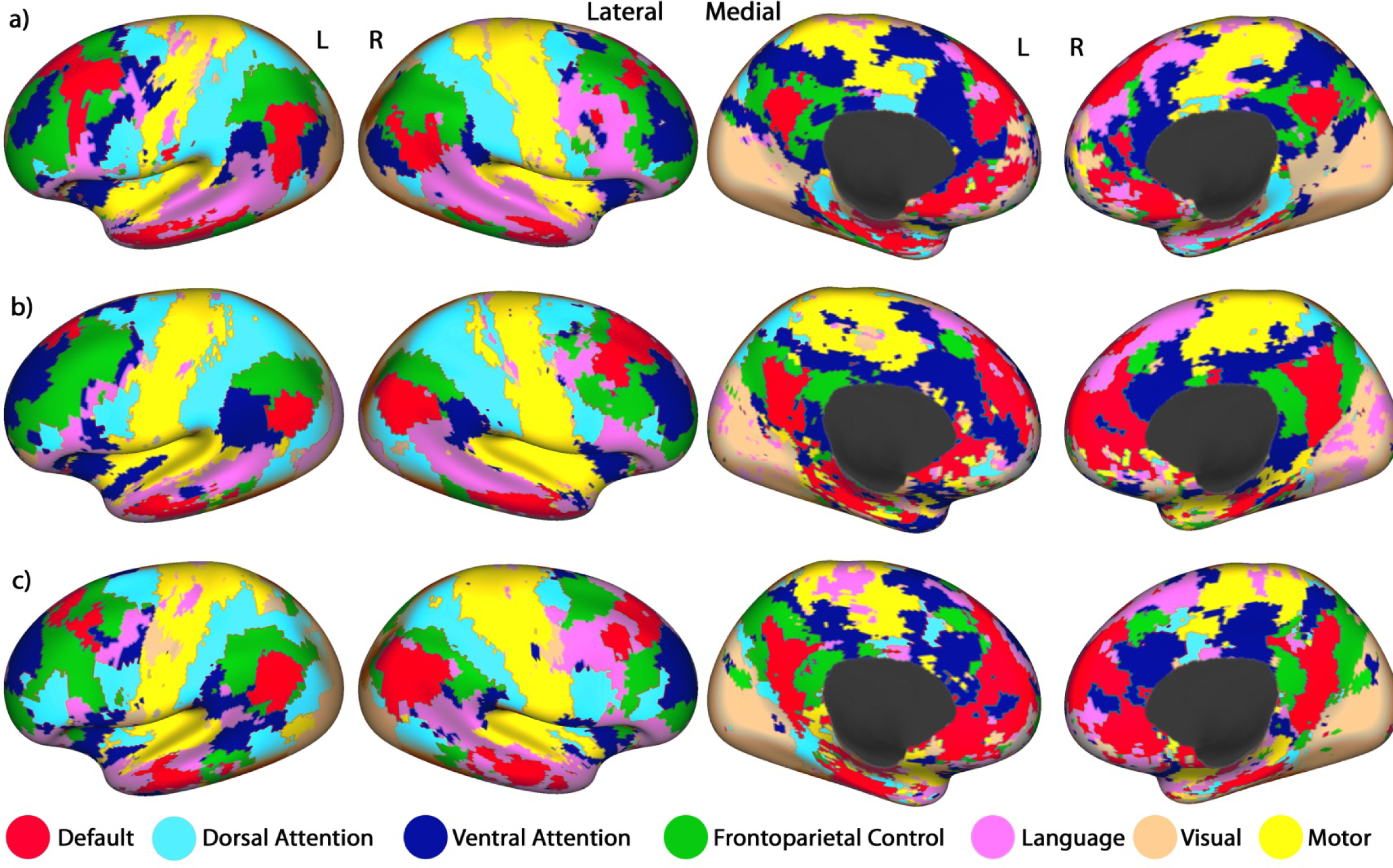
**Winner-take-all maps of individualized resting-state network boundaries** for (a) the experimental subject, (b) group mean of healthy controls, and (c) a representative example of a healthy control. For visualization, maps are projected onto a mean inflated surface from the Human Connectome Project – this schematic representation is used rather than the subject’s own brain in order to clearly visualize between-subject differences. The subject’s individualized default, dorsal attention, and ventral attention parcels (a) were utilized for seed-based connectivity analysis.

Treatment-induced changes were determined using seed-based FC analysis between pre-defined seed pairs, including DAN to DMN, sgACC to DMN, medial orbitofrontal cortex (mOFC) to nucleus accumbens (NAcc), and MTL to DMN. Seed-based correlation was also assessed between other related regions of interest (table S1). These values were compared between pre-treatment scans, post-treatment scans, and both comparator groups.

The subject also received additional rsfMRI scans before and after the sixth and fifteenth treatments. Treatment-induced changes in connectivity were compared in an exploratory manner in order to investigate potential mechanisms of treatment.

### rTMS treatment

The subject was randomized to receive a double-blind course of 20 active treatments over a one-month period. Each treatment session included 4000 left-sided excitatory pulses and 1000 right-sided inhibitory pulses. The detailed treatment protocol is described in the supplement.

## Results

### Clinical response

MADRS score improved by 72% with treatment and remained at the same level upon 6-week follow-up. Secondary outcome variables, including personality scales, cognitive test scores, and self-report emotion scales, are summarized in table 1. The subject experienced no seizures, headaches, or other persistent adverse effects. Transient twitching of facial muscles occurred during treatment, but this was not associated with pain or persistent discomfort. He also incidentally reported a reduction in nicotine cravings, and successfully discontinued cigarette use over the course of the study.

**Table 1.**
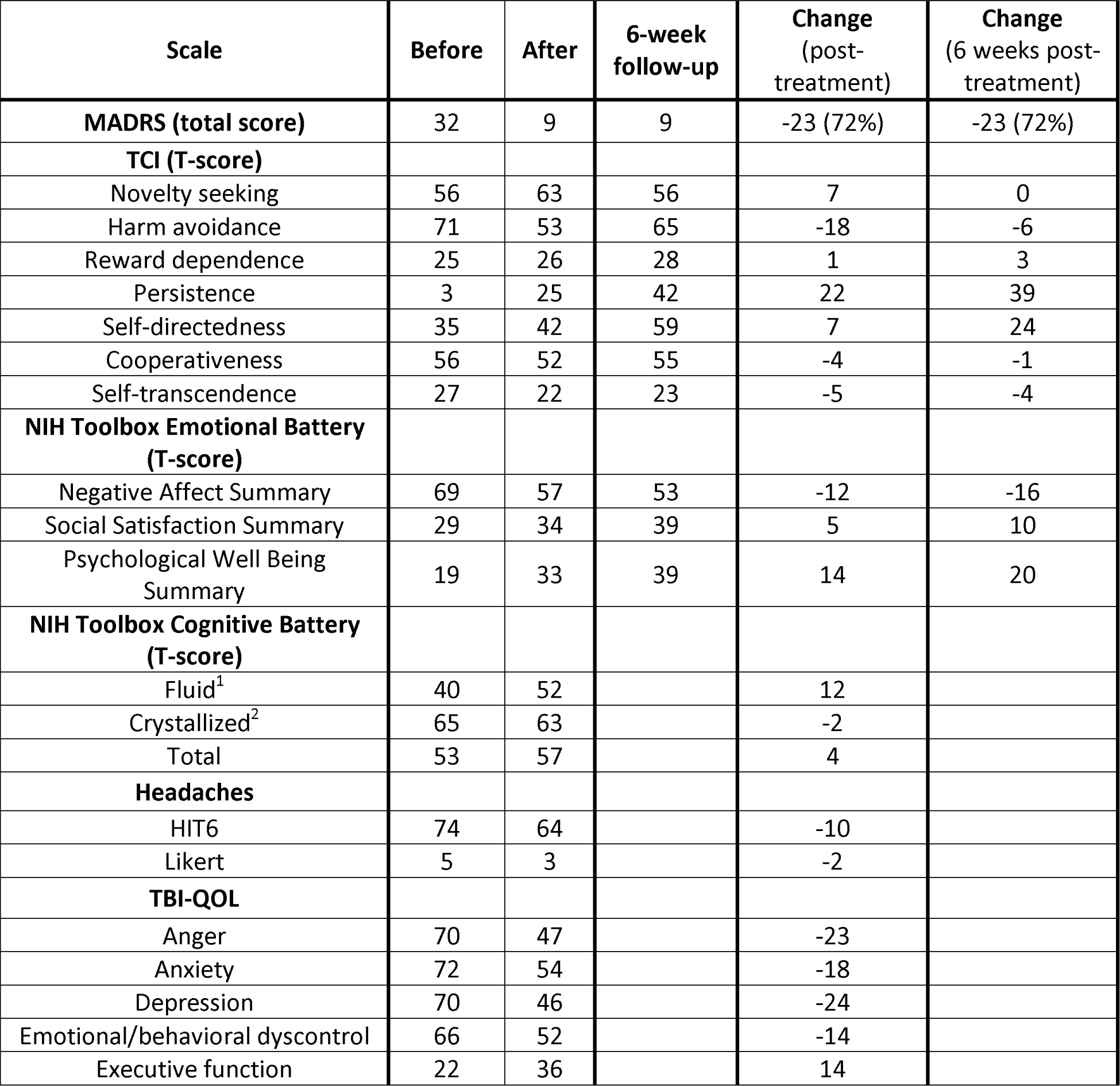
Changes from baseline to 6-week follow-up in MADRS (primary outcome), personality scores, and self-report emotion scores; changes from baseline to post-treatment in cognitive, headache, and TBI-QoL scales.

| Scale | Before | After | 6-week follow-up | Change (post-treatment) | Change (6 weeks post-treatment) |
| --- | --- | --- | --- | --- | --- |
| **MADRS (total score)** | 32 | 9 | 9 | -23 (72%) | -23 (72%) |
| **TCI (T-score)** |  |  |  |  |  |
| Novelty seeking | 56 | 63 | 56 | 7 | 0 |
| Harm avoidance | 71 | 53 | 65 | -18 | -6 |
| Reward dependence | 25 | 26 | 28 | 1 | 3 |
| Persistence | 3 | 25 | 42 | 22 | 39 |
| Self-directedness | 35 | 42 | 59 | 7 | 24 |
| Cooperativeness | 56 | 52 | 55 | -4 | -1 |
| Self-transcendence | 27 | 22 | 23 | -5 | -4 |
| **NIH Toolbox Emotional Battery (T-score)** |  |  |  |  |  |
| Negative Affect Summary | 69 | 57 | 53 | -12 | -16 |
| Social Satisfaction Summary | 29 | 34 | 39 | 5 | 10 |
| Psychological Well Being Summary | 19 | 33 | 39 | 14 | 20 |
| **NIH Toolbox Cognitive Battery (T-score)** |  |  |  |  |  |
| Fluid<sup>1</sup> | 40 | 52 |  | 12 |  |
| Crystallized<sup>2</sup> | 65 | 63 |  | -2 |  |
| Total | 53 | 57 |  | 4 |  |
| **Headaches** |  |  |  |  |  |
| HIT6 | 74 | 64 |  | -10 |  |
| Likert | 5 | 3 |  | -2 |  |
| **TBI-QOL** |  |  |  |  |  |
| Anger | 70 | 47 |  | -23 |  |
| Anxiety | 72 | 54 |  | -18 |  |
| Depression | 70 | 46 |  | -24 |  |
| Emotional/behavioral dyscontrol | 66 | 52 |  | -14 |  |
| Executive function | 22 | 36 |  | 14 |  |

1 Fluid cognition refers to functions unrelated to prior education/experience; in the NIH Toolbox, this includes testing of attention, executive function, episodic memory, working memory, and processing speed.

2 Crystallized cognition refers to functions that involve retrieval of long-term memory; in the NIH Toolbox, this includes testing of reading and vocabulary.

### Identification of novel treatment targets

The individualized map of DAN-DMN anticorrelation (figure 1) identified targets that were anatomically distinct from those identified by previously-reported methods (figure 3). This included comparison with a consensus anatomical MRI-based treatment site, the standard clinical “5 cm rule” site (identified as a location 5 cm anterior to a site on the motor cortex where stimulation induces a contraction in the contralateral abductor pollicis brevis muscle), and a site identified based on peak rsfMRI anticorrelation with a group-mean coordinate for sgACC (described further in supplemental methods).

**Figure 3.**
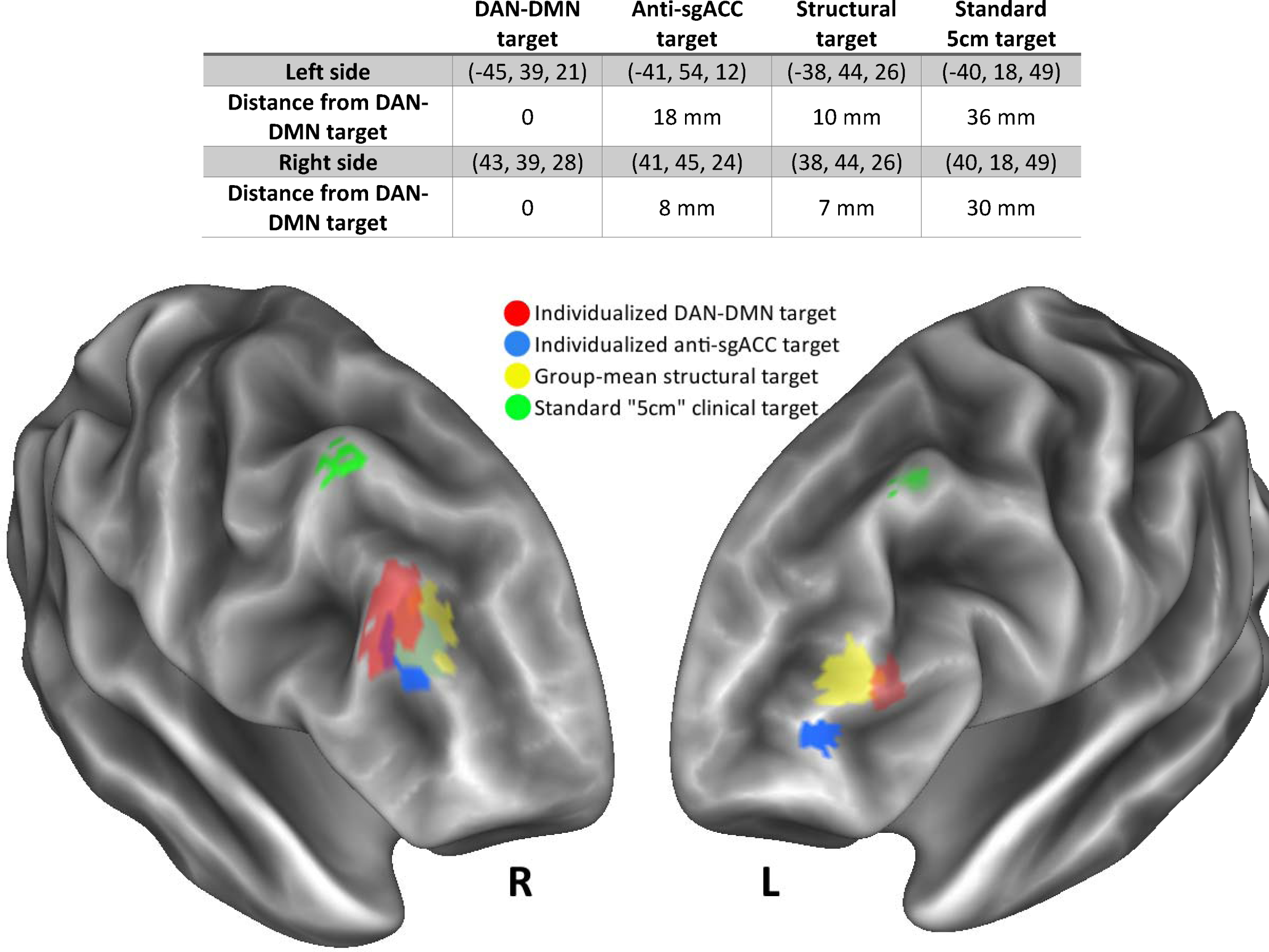
**Anatomical locations of targets** generated by using individualized RSN mapping, individualized sgACC anticorrelation, group-mean structural targeting, and traditional clinical “5-cm rule” target. Colored patches represent an estimated TMS stimulation volume based on spatial distribution of cortical regions within 15 mm of each target (projected onto a surface reconstruction generated from the subject’s anatomical T1-weighted scan). Listed coordinates are in common Talairach atlas space.

In the right hemisphere, there was some overlap between the expected stimulation volumes from the three imaging-based targeting approaches, but a spatial distinction remained evident. In the left hemisphere, there was minimal overlap between the different target volumes. The spatial distance between DAN-DMN targets and the other imaging-based targets ranged between 7 mm and 18 mm (figure 3). This was quite distinct from the clinical “5 cm” rule target sites, which were at least 30 mm away on both sides. The effects of stimulation have been shown to extend approximately 12-16 mm from the stimulation site based on language mapping experiments^41^ and functional connectivity analyses^42^. This suggests that our novel targeting paradigm may be clinically distinct from prior approaches.

### Resting-state functional connectivity changes with treatment

Seed-based correlation analysis suggested differences between the active subject and both comparator groups in mood/reward circuit correlations and other large-scale network correlations (figure 4). Medial temporal limbic regions (MTL) showed higher correlation with DMN in comparison to healthy controls despite lower MTL-DMN connectivity in other subjects with TBI-associated depression; after treatment, this correlation decreased to a level that was comparable to that of healthy controls (Figure 4, left panel). DAN to DMN correlation in the active subject was higher than both control groups and increased further with treatment (Figure 4, right panel). Subgenual anterior cingulate cortex (sgACC) to DMN connectivity was higher than all healthy controls and most TBI-associated depression subjects, while medial orbitofrontal cortex (mOFC) to nucleus accumbens (NAcc) connectivity showed the converse; after treatment, both of these values were comparable to healthy controls (Figure 4, center panels). Additional connectivity changes between these and other related regions are described in the supplement (figure S4). Statistical testing was not conducted on these values because the subject was part of an ongoing clinical trial.

**Figure 4.**
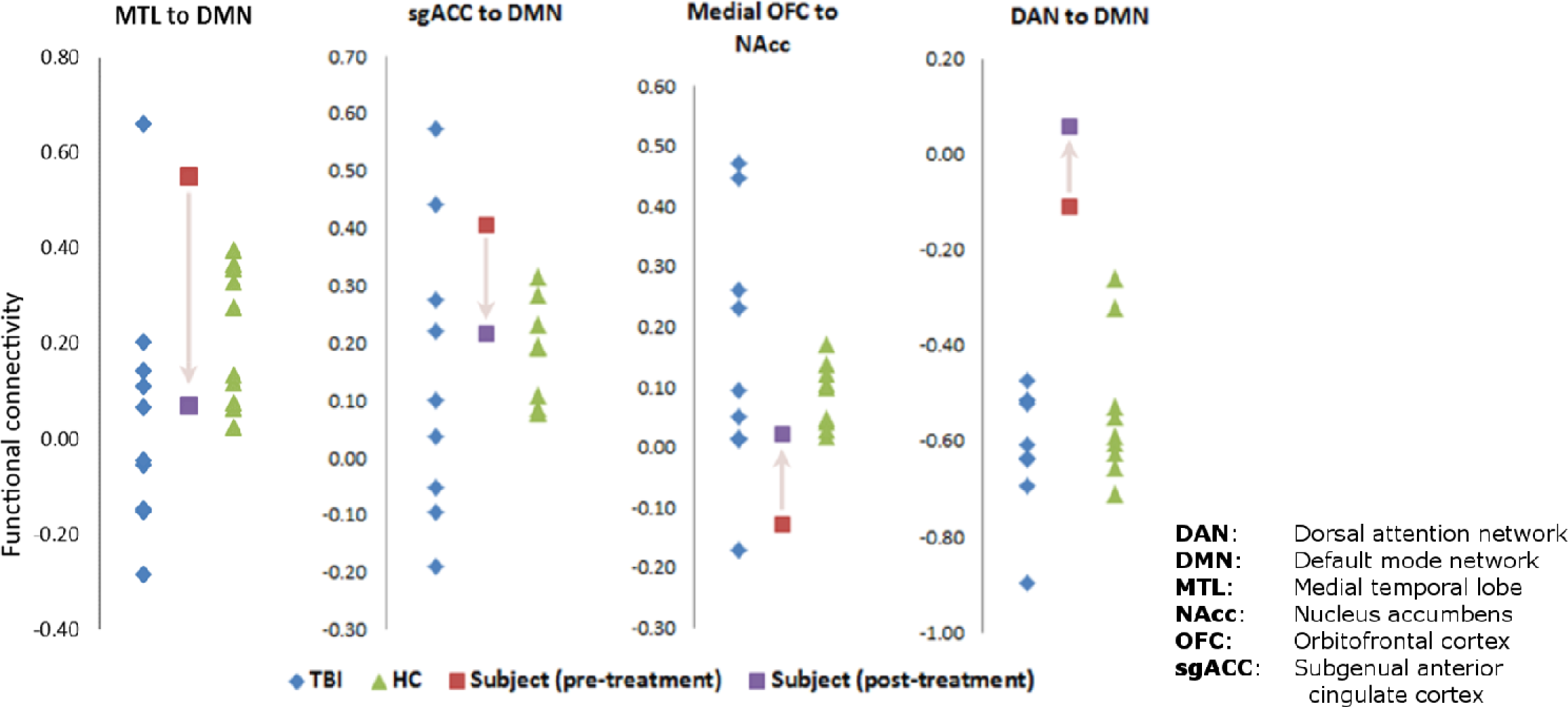
Seed-based connectivity before and after treatment. Comparison of baseline resting-state functional connectivity between specific large-scale networks and cortico-limbic-striatal reward circuits for active subject (before and after treatment), control subjects with TBI-associated depression, and healthy controls.

Qualitatively, whole-brain seed-to-voxel connectivity maps showed that anticorrelation between the left-sided stimulation site and bilateral DMN nodes (ventromedial/dorsomedial prefrontal cortices, precuneus, and temporal poles) was attenuated with treatment (figure 5a). The right-sided stimulation site appeared to show a weaker pattern of anti-correlation with DMN (figure 5b). Treatment-induced changes in connectivity with both stimulation sites appeared to overlap closely with the subject’s individualized DMN parcellation (figure 5), although this effect was again more prominent for the leftsided target.

**Figure 5.**
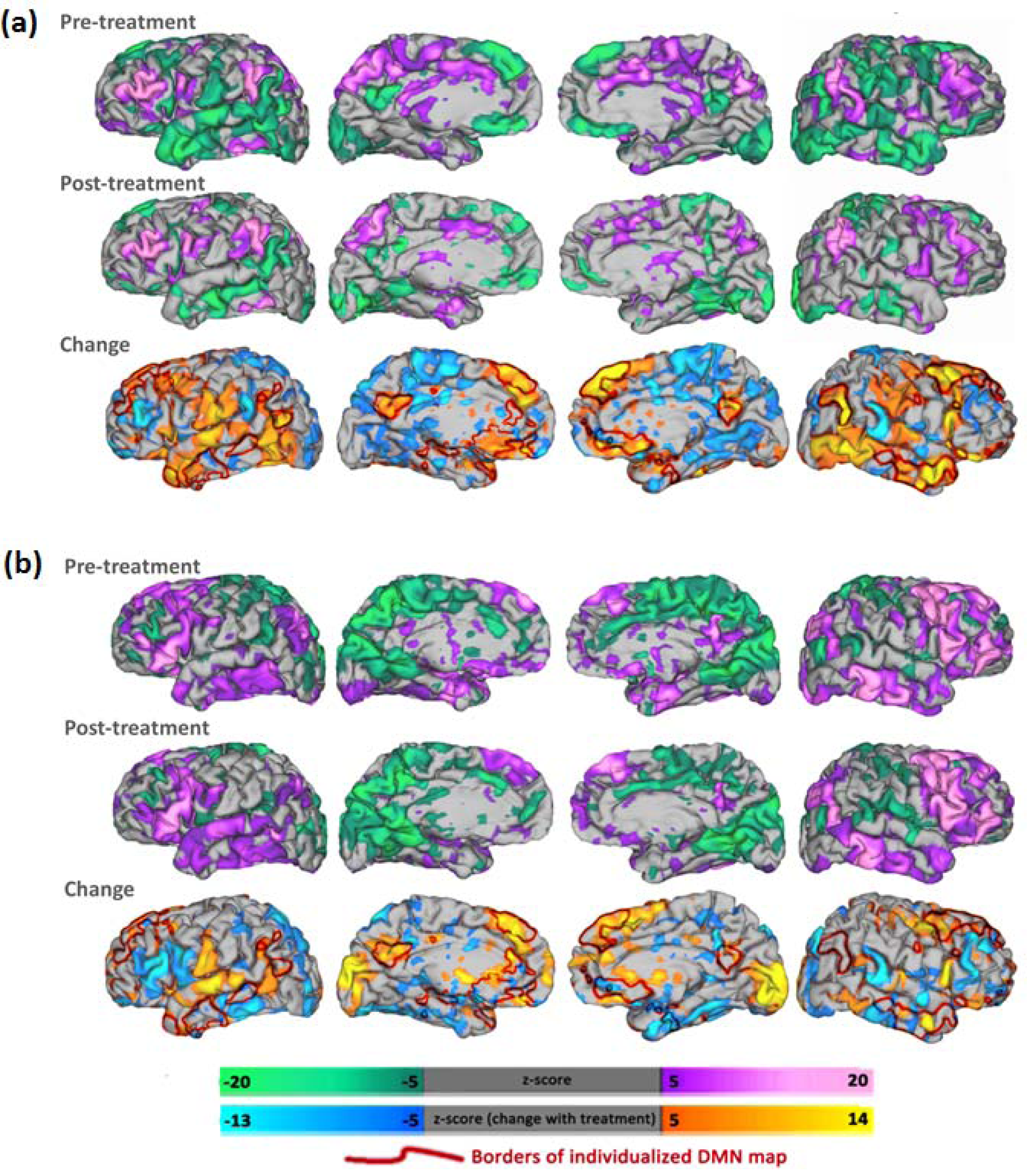
Whole-brain connectivity before and after treatment. (a) Seed-based correlation with a 15-mm spherical ROI at the left-sided rTMS target, including pre-treatment map (top), post-treatment map (middle), and change with treatment (bottom) depicted along with individualized DMN borders (red lines). (b) Seed-based correlation with right-sided rTMS target, including pre-treatment map (top), posttreatment map (middle), and change with treatment (bottom) depicted along with individualized DMN borders (red)

### Mid-treatment changes

Additional scans collected during the treatment course showed that the treatment successfully attenuated anti-correlation between the stimulation site and sgACC (figure 6a). Target-sgACC anticorrelation progressively decreased over the course of the four longitudinal time points. The acute effects of treatment, meanwhile, led to substantial increases in target-sgACC anti-correlation, suggesting that the longitudinal increase may be mediated by a compensatory homeostatic response.

**Figure 5.**
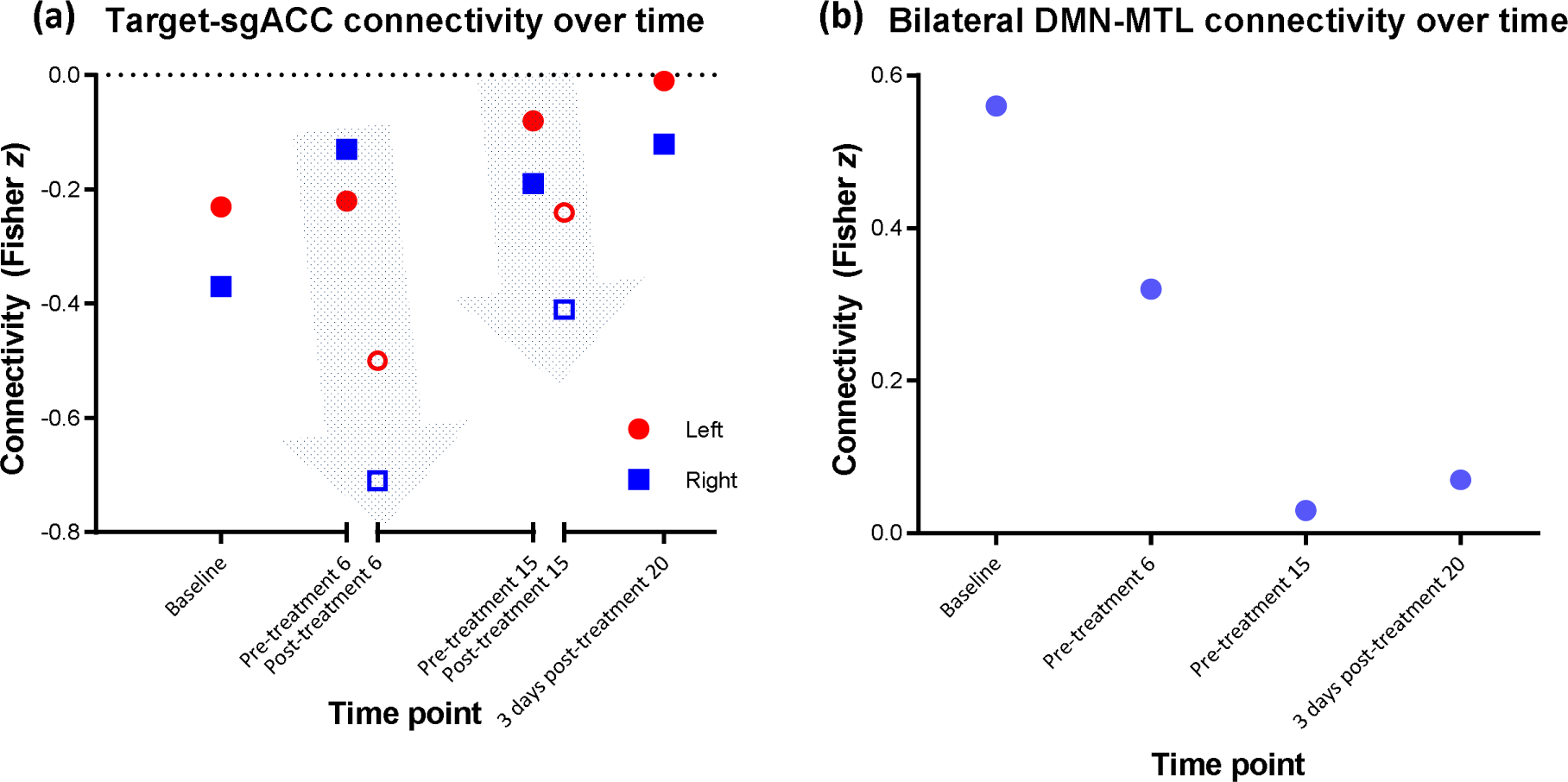
Changes in connectivity over time. (a) Individual treatments led to substantial increase in anticorrelation between each stimulation site (left/right) and sgACC; gray arrows depict the change between pre-treatment and post-treatment scans over the course of a single session. The longitudinal treatment course led to a gradual attenuation of this anticorrelation. (b) Bilateral DMN-MTL connectivity, which was substantially elevated at baseline, also showed a gradual decline over the course of treatment.

Given the striking magnitude of the elevation in baseline MTL-DMN connectivity and normalization with treatment, it was unclear whether this may be attributable to regression to the mean – if the baseline measurement was an extreme, noise-driven outlier, the second measurement would appear more normal simply by chance. However, post-hoc analysis on mid-treatment scans revealed that the normalization of MTL-DMN connectivity occurred gradually over the treatment course and stabilized over the last few treatments (Figure 6b). These mid-treatment changes may serve as a promising biomarker for early prediction of treatment response.

## Discussion

To our knowledge, this is the first report of successful use of rTMS in mood disorders associated with repetitive TBI. Treatment led to improvements in clinician-assessed mood ratings, self-report emotional scores (including ratings of mood, anger, anxiety, and behavioral dyscontrol), and fluid cognition. Quantitative personality testing showed particularly drastic changes in persistence, a measure of reward-based learning which is related to mOFC-NAcc connectivity^43^ and has some predictive value for rTMS response^44^. This was accompanied by apparent normalization of connectivity in circuits related to reward-based learning and reward-motivated behavior, although this is difficult to assess conclusively in a single subject. Adverse effects were limited to transient twitching of facial muscles during treatment. Although rTMS can cause headaches^11^, the subject reported an improvement in baseline headaches.

Our successful use of individualized RSN mapping for rTMS guidance demonstrates the potential for broader clinical applications for resting-state fMRI, which are currently limited to pre-surgical mapping^45^. Notably, the rTMS targets identified in this subject were spatially distinct from targets identified by prior MRI-based approaches. The distance between these targets (7-18 mm) was comparable to the expected radius of stimulation with rTMS^41,42^, suggesting there was likely minimal overlap in the stimulation volumes between different targets. All three MRI-based targets were at least 3 cm away from standard “5 cm” targets, which may be partly explained by the fact that the 5 cm rule does not consider interindividual differences in head/brain size and the subject, unsurprisingly, had a larger-than-average head. While this is consistent with findings that structural MRI guidance improves clinical rTMS outcomes^46^, rsfMRI-guided treatment has not yet been systematically compared with other targeting approaches. The remarkable clinical benefit in this individual warrants further investigation to answer the question of how to optimize rTMS targeting.

While RSN mapping enables subject-specific targeting of specific functional regions, it remains unclear which region should be chosen for optimal clinical outcome. Recent literature suggests that rTMS response is related to baseline DMN correlation with sgACC^12^ as well as group-mean anti-correlation between the DLPFC treatment site and sgACC^18^. Depression in TBI is associated with dysfunction in mood-regulating circuits involving DMN and limbic connectivity with DAN^31,32^, a system that is heavily involved in selection of stimuli based on internal expectations. Repetitive head trauma has additionally been associated with pathological and neuroimaging changes in prefrontal and medial temporal limbic regions^36,37,47^, which may explain the success of a structurally-oriented treatment despite failure of traditional pharmacotherapy. These early findings informed the theoretical underpinnings of our targeting approach, but further research is necessary to investigate the clinical utility of other network targets identified via individualized mapping.

The gradual changes in target-sgACC anti-correlation and MTL-DMN correlation may both serve as promising biomarkers for early prediction of treatment response. Individual stimulation sessions led to substantial increase in target-sgACC anti-correlation, while the longitudinal treatment course led to overal attenuation of this value; this may be mediated by a compensatory homeostatic response. DLPFC-sgACC anti-correlation is a cornerstone of many neurophysiologic models of major depression, but it has been difficult to trace longitudinally in an individual subject without individualizing the optimal DLPFC region. Similarly, MTL-DMN connectivity has been implicated in chronic traumatic encephalopathy, but has also been difficult to trace individually without individualizing the DMN maps.

This study also raises questions regarding resting-state functional connectivity differences between single and repetitive TBI. The experimental subject showed notable baseline differences in comparison with both healthy controls and individuals with single TBI-associated depression. The most striking difference appeared to be in connectivity between medial temporal lobe and the subject-specific DMN map, which demonstrated abnormalities in the opposite direction of the changes found in patients with single TBI-associated depression. The aberrant MTL connectivity observed here is consistent with emerging findings implicating MTL as a common site of pathology in chronic traumatic encephalopathy (CTE), a neurodegenerative tauopathy associated with repetitive head trauma in athletes^36,37,47^. Given the lack of a validated method for detection of CTE in a living person, further investigation should aim to elucidate the relationship between CTE-related tau pathology, functional connectivity, and mood disorders.

Limitations of this study include the inability to assess the impact of intensity, frequency, duration, or laterality of rTMS treatment. The use of adjunctive right-sided low-frequency stimulation, which is believed to be effective for anxiety disorders and post-traumatic stress disorder (PTSD)^48,49^, may have contributed to the observed clinical improvement, as the neuropsychiatric sequelae of repetitive concussive TBI include a mixture of depression, anxiety, and behavioral disinhibition rather than a single categorical mood disorder. We also administered a relatively high treatment dose which may have contributed to the subject’s improvement.

We are unable to clearly disentangle the observed clinical response from placebo effect in this individual subject. Placebo effect appears somewhat less likely given the stability of improvement upon 6-week post-treatment follow-up, although the long-term durability of treatment effects and the potential need for maintenance treatments remain unclear. Placebo effect also appears less likely given the observation of gradual changes in DLPFC-sgACC anti-correlation which showed consistent patterns of both acute and longitudinal change. Furthermore, the gradual normalization of MTL-DMN connectivity suggests possible modulation of regions that are heavily implicated in postmortem studies of CTE.

In order to address some of these limitations, our future work will include completion of the ongoing randomized-controlled trial in a larger sample of patients with mood disorders, anxiety disorders, and PTSD associated with TBI. We are also prospectively investigating the effect of precise stimulation site on changes in network connectivity in prefrontal cortex. This lays the foundation for development of personalized neurostimulation based on patient-specific brain network disturbances not only for patients with repetitive TBI and high risk for CTE, but also potentially for any neuropsychiatric condition with identifiable network dysfunction.

## Acknowledgements

We would especially like to thank the experimental participant as well as the participants in the comparator groups. We thank Drs. Eric Leuthardt and Abraham Snyder for logistical support with implementation of individualized RSN mapping. We thank Xin Hong and Linda Hood for technical support with rTMS and MRI equipment. We thank Drs. Sindhu Jacob and Martin Wice for referring participants. We acknowledge Drs. Bradley Schlaggar, Charles Zorumski, and David Soleimani-Meigooni for critically reviewing the manuscript. We additionally acknowledge valuable discussions with Drs. Michael Fox, Steven Petersen, Charles Conway, Eric Wasserman, Sarah Lisanby, Bruce Luber, Andrew Drysdale, Irving Reti, Vani Rao, Maurizio Corbetta, Gordon Shulman, and Abraham Snyder which led to the conception and refinement of the study design. The first author would additionally like to thank Drs. Kevin Black, Pilar Cristancho, Charles Zorumski, and C. Robert Cloninger for extensive longitudinal guidance leading to the conception of this study.

## Author contributions

SHS, DLB, ARC, and NTT designed the clinical protocol. ARC provided rTMS expertise and equipment as well as functional connectivity expertise. CDH and TOL provided expertise and novel analytical scripts for individualized RSN mapping and functional connectivity analysis. SK provided expertise and developed scripts for functional connectivity processing. SHS coordinated the study, recruited subjects, administered treatments, processed MRI data, and conducted functional connectivity analyses. SHS, NTT, and PS acquired MRI scans and conducted clinical assessments. SHS and DLB wrote the manuscript with intellectual contributions from all authors.

## Additional funding information

This study was funded by pilot funds from the McDonnell Center for Systems Neuroscience and the Mallinckrodt Institute of Radiology at Washington University.

Data were provided in part by the Human Connectome Project, WU-Minn Consortium (Principal Investigators: David Van Essen and Kamil Ugurbil; 1U54MH091657) funded by the 16 NIH Institutes and Centers that support the NIH Blueprint for Neuroscience Research; and by the McDonnell Center for Systems Neuroscience at Washington University.

## References

1. Fox MD, Liu H, Pascual-Leone A. Identification of reproducible individualized targets for treatment of depression with TMS based on intrinsic connectivity. NeuroImage. 2013;66:151-160.

2. Fox MD, Buckner RL, Liu H, Chakravarty MM, Lozano AM, Pascual-Leone A. Resting-state networks link invasive and noninvasive brain stimulation across diverse psychiatric and neurological diseases. Proceedings of the National Academy of Sciences of the United States of America. 2014;111(41):E4367-4375.

3. Klomjai W, Lackmy-Vallee A, Roche N, Pradat-Diehl P, Marchand-Pauvert V, Katz R. Repetitive transcranial magnetic stimulation and transcranial direct current stimulation in motor rehabilitation after stroke: An update. Ann Phys Rehabil Med. 2015;58(4):220-224.

4. Pascual-Leone A, Tormos JM, Keenan J, Tarazona F, Canete C, Catala MD. Study and modulation of human cortical excitability with transcranial magnetic stimulation. J Clin Neurophysiol. 1998;15(4):333-343.

5. Fecteau S, Dickler M, Pelayo R, et al. Cortical Excitability During Passive Action Observation in Hospitalized Adults With Subacute Moderate to Severe Traumatic Brain Injury: A Preliminary TMS Study. Neurorehabil Neural Repair. 2015;29(6):548-556.

6. Reti IM, Schwarz N, Bower A, Tibbs M, Rao V. Transcranial magnetic stimulation: A potential new treatment for depression associated with traumatic brain injury. Brain injury: [BI]. 2015;29(7-8):789-797.

7. Dinan TG, Mobayed M. Treatment resistance of depression after head injury: a preliminary study of amitriptyline response. Acta Psychiatr Scand. 1992;85(4):292-294.

8. Fann JR, Bombardier CH, Temkin N, et al. Sertraline for Major Depression During the Year Following Traumatic Brain Injury: A Randomized Controlled Trial. J Head Trauma Rehabil. 2017;32(5):332-342.

9. Pitkanen A, Immonen R. Epilepsy related to traumatic brain injury. Neurotherapeutics. 2014;11(2):286-296.

10. Scheid R, von Cramon DY. Clinical findings in the chronic phase of traumatic brain injury: data from 12 years’ experience in the Cognitive Neurology Outpatient Clinic at the University of Leipzig. Dtsch Arztebl Int. 2010;107(12):199-205.

11. Perera T, George MS, Grammer G, Janicak PG, Pascual-Leone A, Wirecki TS. The Clinical TMS Society Consensus Review and Treatment Recommendations for TMS Therapy for Major Depressive Disorder. Brain stimulation.9(3):336-346.

12. Liston C, Chen AC, Zebley BD, et al. Default mode network mechanisms of transcranial magnetic stimulation in depression. Biological psychiatry. 2014;76(7):517-526.

13. Dougherty DD, Weiss AP, Cosgrove GR, et al. Cerebral metabolic correlates as potential predictors of response to anterior cingulotomy for treatment of major depression. J Neurosurg. 2003;99(6):1010-1017.

14. Mayberg HS, Liotti M, Brannan SK, et al. Reciprocal limbic-cortical function and negative mood: converging PET findings in depression and normal sadness. Am J Psychiatry. 1999;156(5):675-682.

15. Drevets WC, Bogers W, Raichle ME. Functional anatomical correlates of antidepressant drug treatment assessed using PET measures of regional glucose metabolism. Eur Neuropsychopharmacol. 2002;12(6):527-544.

16. Nobler MS, Oquendo MA, Kegeles LS, et al. Decreased regional brain metabolism after ect. Am J Psychiatry. 2001;158(2):305-308.

17. Mayberg HS, Lozano AM, Voon V, et al. Deep brain stimulation for treatment-resistant depression. Neuron. 2005;45(5):651-660.

18. Fox MD, Buckner RL, White MP, Greicius MD, Pascual-Leone A. Efficacy of transcranial magnetic stimulation targets for depression is related to intrinsic functional connectivity with the subgenual cingulate. Biological psychiatry. 2012;72(7):595-603.

19. Siddiqi SH, Weigand AT, Pascual-Leone A, Fox MD. Individualized connectivity between rTMS targets and the subgenual cingulate is unrelated to antidepressant response Paper presented at: 2nd European Conference on Brain Stimulation in Psychiatry2017; Munich.

20. Gordon EM, Laumann TO, Adeyemo B, et al. Individual-specific features of brain systems identified with resting state functional correlations. NeuroImage. 2016.

21. Gordon EM, Laumann TO, Adeyemo B, Petersen SE. Individual Variability of the System-Level Organization of the Human Brain. Cerebral cortex (New York, NY: 1991). 2015.

22. Hacker CD, Laumann TO, Szrama NP, et al. Resting state network estimation in individual subjects. Neuroimage. 2013;82:616-633.

23. Laumann TO, Gordon EM, Adeyemo B, et al. Functional System and Areal Organization of a Highly Sampled Individual Human Brain. Neuron. 2015;87(3):657-670.

24. Wang D, Buckner RL, Fox MD, et al. Parcellating cortical functional networks in individuals. Nature neuroscience. 2015;18(12):1853-1860.

25. Glasser MF, Coalson TS, Robinson EC, et al. A multi-modal parcellation of human cerebral cortex. Nature. 2016;536(7615):171-178.

26. Gordon EM, Laumann TO, Adeyemo B, Huckins JF, Kelley WM, Petersen SE. Generation and Evaluation of a Cortical Area Parcellation from Resting-State Correlations. Cerebral cortex (New York, NY: 1991). 2016;26(1):288-303.

27. Kaiser RH, Andrews-Hanna JR, Wager TD, Pizzagalli DA. Large-Scale Network Dysfunction in Major Depressive Disorder: A Meta-analysis of Resting-State Functional Connectivity. JAMA psychiatry. 2015;72(6):603-611.

28. Choi KS, Riva-Posse P, Gross RE, Mayberg HS. Mapping the “Depression Switch” During Intraoperative Testing of Subcallosal Cingulate Deep Brain Stimulation. JAMA neurology. 2015;72(11):1252-1260.

29. Mac Donald CL, Johnson AM, Cooper D, et al. Detection of blast-related traumatic brain injury in U.S. military personnel. The New England journal of medicine. 2011;364(22):2091-2100.

30. Han K, Mac Donald CL, Johnson AM, et al. Disrupted modular organization of resting-state cortical functional connectivity in U.S. military personnel following concussive ‘mild’ blast‐related traumatic brain injury. NeuroImage. 2014;84:76-96.

31. Han K, Chapman SB, Krawczyk DC. Altered Amygdala Connectivity in Individuals with Chronic Traumatic Brain Injury and Comorbid Depressive Symptoms. Front Neurol. 2015;6:231-231.

32. Han K, Chapman SB, Krawczyk DC. Disrupted Intrinsic Connectivity among Default, Dorsal Attention, and Frontoparietal Control Networks in Individuals with Chronic Traumatic Brain Injury. Journal of the International Neuropsychological Society: JINS. 2016;22(2):263-279.

33. van der Horn HJ, Liemburg EJ, Scheenen ME, de Koning ME, Spikman JM, van der Naalt J. Graph Analysis of Functional Brain Networks in Patients with Mild Traumatic Brain Injury. PLoS One. 2017;12(1):e0171031-e0171031.

34. Caeyenberghs K, Verhelst H, Clemente A, Wilson PH. Mapping the functional connectome in traumatic brain injury: What can graph metrics tell us? Neuroimage. 2016.

35. Sharp DJ, Scott G, Leech R. Network dysfunction after traumatic brain injury. Nature reviews Neurology. 2014;10(3):156-166.

36. Coughlin JM, Wang Y, Minn I, et al. Imaging of Glial Cell Activation and White Matter Integrity in Brains of Active and Recently Retired National Football League Players. JAMA neurology. 2017;74(1):67-74.

37. Stein TD, Alvarez VE, McKee AC. Chronic traumatic encephalopathy: a spectrum of neuropathological changes following repetitive brain trauma in athletes and military personnel. Alzheimer’s research & therapy. 2014;6(1):4.

38. Montenigro PH, Alosco ML, Martin BM, et al. Cumulative Head Impact Exposure Predicts Later‐Life Depression, Apathy, Executive Dysfunction, and Cognitive Impairment in Former High School and College Football Players. Journal of neurotrauma. 2017;34(2):328-340.

39. Van Essen DC, Ugurbil K, Auerbach E, et al. The Human Connectome Project: a data acquisition perspective. NeuroImage. 2012;62(4):2222-2231.

40. Power JD, Mitra A, Laumann TO, Snyder AZ, Schlaggar BL, Petersen SE. Methods to detect, characterize, and remove motion artifact in resting state fMRI. NeuroImage. 2014;84:320-341.

41. Sollmann N, Hauck T, Tussis L, et al. Results on the spatial resolution of repetitive transcranial magnetic stimulation for cortical language mapping during object naming in healthy subjects. BMC Neurosci. 2016;17(1):67.

42. Opitz A, Fox MD, Craddock RC, Colcombe S, Milham MP. An integrated framework for targeting functional networks via transcranial magnetic stimulation. NeuroImage. 2016;127:86-96.

43. Gusnard DA, Ollinger JM, Shulman GL, et al. Persistence and brain circuitry. Proc Natl Acad Sci U S A. 2003;100(6):3479-3484.

44. Siddiqi SH, Chockalingam R, Cloninger CR, Lenze EJ, Cristancho P. Use of the Temperament and Character Inventory to Predict Response to Repetitive Transcranial Magnetic Stimulation for Major Depression. Journal of psychiatric practice. 2016;22(3):193-202.

45. Lee MH, Miller-Thomas MM, Benzinger TL, et al. Clinical Resting-state fMRI in the Preoperative Setting: Are We Ready for Prime Time? Top Magn Reson Imaging. 2016;25(1):11-18.

46. Fitzgerald PB, Hoy K, McQueen S, et al. A randomized trial of rTMS targeted with MRI based neuro-navigation in treatment-resistant depression. Neuropsychopharmacology: official publication of the American College of Neuropsychopharmacology. 2009;34(5):1255-1262.

47. Coughlin JM, Wang Y, Munro CA, et al. Neuroinflammation and brain atrophy in former NFL players: An in vivo multimodal imaging pilot study. Neurobiol Dis. 2015;74:58-65.

48. Diefenbach GJ, Bragdon LB, Zertuche L, et al. Repetitive transcranial magnetic stimulation for generalised anxiety disorder: a pilot randomised, double-blind, sham-controlled trial. Br J Psychiatry. 2016;209(3):222-228.

49. Wassermann EM, Zimmermann T. Transcranial magnetic brain stimulation: therapeutic promises and scientific gaps. Pharmacology & therapeutics. 2012;133(1):98-107.

